# Independent evolutionary trajectories of genomic repeats and non–repeat genome features in Actinomycetota

**DOI:** 10.64898/2026.06.01.729349

**Authors:** Chandana Gobattini, Uday Kumar Ammisetty, Bhargav Reddy Konkala, Akash Ajay

## Abstract

Genomic repeats, particularly simple sequence repeats, influence genome stability and gene regulation. While many genomic traits exhibit phylogenetic signal and pulsed evolution, repeat elements have rarely been examined in a comparative framework, and their evolutionary relationships with other genome features remain poorly understood. We contrasted the trait evolution of genomic repeats with non–repeat traits across diverse actinobacterial orders, testing for phylogenetic signal, evolutionary mode, and pulsed dynamics using time–calibrated, 16S rRNA, and whole–genome sequence (WGS) trees. Non–repeat traits consistently exhibited signs of punctuated evolution, with larger pulses in species–rich orders such as *Mycobacteriales* and Actinomycetales; repeats, by contrast, were evolutionarily labile, lacking phylogenetic signal and largely decoupled from other genomic traits. Bifidobacteriales was the sole exception, with repeats exhibiting phylogenetic signal only under the WGS tree. The WGS tree also recovered a stronger signal for genome–level traits, highlighting its utility in comparative analyses. Genome size evolution in Actinomycetota appears driven primarily by protein–coding gene expansion rather than repeat accumulation.

## 1. Introduction

Genomic repeats are ubiquitous across the tree of life and exert far–reaching influences on genome architecture, including elevated local mutation rates (Byzmek et al., 2001; Chen et al, 2009; Lenz et al., 2014; Ajay et al., 2024), genome instability (Byzmek et al., 2001; McGinty et al., 2025), and modulation of gene expression (Liao et al., 2023). Broadly, repetitive sequences fall into two major classes: interspersed repeats, which include transposable elements and retrotransposons, and tandem repeats, which encompass satellite DNA, minisatellites, and microsatellites (Jurka et al., 2007; Richard et al., 2008). Simple sequence repeats (SSRs) are tandem arrays of 1–6 bp motifs (Ahmad et al., 2018) that are hypermutable due to polymerase slippage during replication (Byzmek et al., 2001) and serve as a major source of functional genetic diversity, particularly in bacterial pathogens (Van Belkum, 1999). In prokaryotes, additional repeat types such as CRISPR arrays (Horvath & Barrangou, 2010), insertion sequences (Mahillon & Chandler, 1998; Siguer et al., 2014), and miniature inverted–repeat transposable elements (MITEs) (Lu et al., 2012) further contribute to genome plasticity through recombination, horizontal gene transfer, and adaptive immunity.

Genomic traits have been increasingly studied within a phylogenetic comparative framework to address questions of genome size evolution, complexity, and adaptation. Such analyses have revealed that traits like genome size, GC content, and gene counts often carry significant phylogenetic signal (Martinez–Gutierrez & Aylward, 2022; Kyriacou et al., 2024) and exhibit pulsed, rather than gradual, evolutionary dynamics (Gao & Wu, 2022; Ajay et al., 2025), with pulse magnitude sometimes correlated with species richness (Ajay et al., 2025). Genome size itself has been implicated in trait plasticity and adaptive radiation (Bhadra et al., 2023). In the case of genomic repeats, a recent study noted that repeat length distributions remain remarkably stable across evolutionary timescales, with longer repeats balancing expansion and contraction processes (McGinty et al., 2025).

While repetitive sequences constitute a substantial fraction of many eukaryotic genomes and are recognized as major drivers of genome size expansion in plants and mammals (Lopez-Flores and Garrido-Romes, 2012; Biscotti et al., 2015), bacterial genomes are generally under strong selection for compactness, maintaining high coding density and relatively few intergenic or repetitive elements (Haubold and Weiehe, 2006; Sela et al., 2016; Bobay and Ochman, 2017). Yet, genomic repeats are not absent from prokaryotes, and the degree to which their abundance, composition, and evolutionary dynamics align with those of other genomic traits—or reflect deeper phylogenetic constraints—remains poorly understood. Prokaryotic repeat elements span a striking structural diversity—from miniature inverted–repeat transposable elements to CRISPR arrays—and serve multifaceted roles in gene regulation, host–pathogen interactions, adaptive immunity, and genome plasticity, making their evolutionary dynamics central to understanding bacterial genome evolution (Delihas, 2011; Chen et al., 2012). Actinomycetota (Actinobacteria) is one of the largest and most ecologically diverse bacterial phyla and provides an appropriate system for examining the macroevolutionary dynamics of genomic traits. To date, the evolution of repeat elements has not been examined within a phylogenetic comparative framework that explicitly tests for phylogenetic signal and pulsed dynamics in any bacterial lineage, apart from a single preliminary study in *Staphylococci* (Ajay et al., 2023), which reported no phylogenetic signal in repeat fraction using whole–genome trees. That finding left unresolved whether the absence of signal reflected limited taxonomic scope, sensitivity to tree topology, or reliance on a single aggregated repeat measure. The present study addresses these questions directly by applying a multi–phylogeny, multi–trait comparative framework to a broad set of actinobacterial orders. To distinguish among these possibilities, the present study expands the phylogenetic comparative analysis and pulsed evolution framework to a broader taxonomic scale (order level), three independent phylogenetic hypotheses—time–calibrated tree (e.g., Wang & Obbard, 2023), 16S rRNA gene tree (e.g., Martiny et al., 2013), and whole–genome sequence tree (e.g., Ajay et al., 2024)—and a richer set of repeat–related traits, including GC–poor and GC–rich repeat counts. This study, therefore, provides the first systematic examination of the phylogenetic signal, tempo, and genomic correlates of repeat elements across multiple actinobacterial orders, employing three independent phylogenetic hypotheses to assess the robustness of these relationships.

## 2. Methodology

### 2.1 Data collection

Data for genomic features - genome size, genomic GC, Number of genes (gene count), number of protein-coding genes, average protein length, average gene length, genes with plus and minus orientation for Actinomycetota were collected from the NCBI assembly database (Kitts et al., 2016). The same genome assemblies were analysed for tandem repeat content using the Galaxy implementation of the Tandem Repeat Finder (*etandem*) (Benson, 1999), run with a minimum repeat unit length of 10 bp and a maximum of 500 bp. The resulting per–genome repeat output was parsed and summarized in R using custom scripts to generate the repeat tables for downstream analyses. The tandem repeats were further classified into distinct classes based on the length of their repeat unit, as shown in Table 1. Furthermore, repeats were classified as GC–rich (> 55 % GC) or GC–poor (< 45 % GC) to examine whether repeat propagation across lineages is biased toward a particular base composition. Taxonomic assignments for all genomes were verified against the NCBI Taxonomy database (Federhen, 2012; Schoch et al., 2020). Table 1 lists the genomic features for which data were collected for Actinomycetota for the following orders: Mycobacteriales, Frankiales, Catenulisporales, Glycomycetales, Jiangellales, Actinomycetales, Bifidobacteriales, Nakamuralles, and Geodermatophilase. The complete dataset is provided in Supplementary Table 1 (ST1).

**Table 1.** Genomic traits examined in this study, grouped into non–repeat genomic traits and repeat–related traits.

| Category | Trait |
| --- | --- |
| Non-repeat genomic traits | Genome size |
|  | Gc% |
|  | Gene Count |
|  | Protein Coding Genes |
|  | Average Protein length |
|  | Average gene length |
|  | Number of stress genes |
|  | Number of DNA binding genes |
|  | Number of Pseudogenes |
|  | Number of repeat containing genes |
|  | Number of Genes with Minus orientation |
|  | Number of Genes with Plus orientation |
|  | Number of Aquaporin genes |
| Repeat-related traits | Number of Mono-nucleotide repeats |
|  | Number of Di-nucleotide repeats |
|  | Number of Tri-nucleotide repeats |
|  | Number of Tetra-nucleotide repeats |
|  | Number of 5–10 bp repeat units |
|  | Number of 10–50 bp repeat units |
|  | Number of 50–100 bp repeat units |
|  | Number of >100 bp repeat units |
|  | total length of repeats |
|  | Number of GC_Poor repeats |
|  | Number of GC_Rich repeats |

### 2.2 Phylogenetic framework reconstruction and downstream comparative methods

To assess the robustness of our phylogenetic comparative results and to exclude the possibility that observed trait–phylogeny associations were artefacts of a particular tree topology or branch–length estimation method, all downstream comparative analyses were independently repeated using three alternative phylogenies of the *Actinomycota*: a time–calibrated supertree from TimeTree (Kumar et al., 2022), a maximum–likelihood tree inferred from 16S rRNA gene sequences, and a whole–genome sequence (WGS) phylogeny derived from a core–genome alignment (or multi–locus concatenation). The 16S rRNA gene tree and the whole–genome phylogeny were both obtained via the Type (Strain) Genome Server (TYGS) (Meier–Kolthoff & Göker, 2019), using its automated GBDP–based tree reconstruction and 16S rRNA gene extraction workflows.

The 16S rRNA tree reflects conserved ribosomal evolution and is a standard for bacterial systematics, though it offers limited resolution at shallower nodes. In contrast, the WGS phylogeny provides substantially higher resolution by incorporating hundreds to thousands of orthologous genes distributed across the genome. The TimeTree supertree adds absolute divergence times, enabling biologically meaningful branch–length scaling required for models such as Brownian motion or Ornstein–Uhlenbeck. Topological congruence among the three phylogenetic hypotheses was quantified using the Robinson–Foulds (symmetric difference) distance (Briand et al., 2020) and the quartet distance (absolute and normalised). All trees were pruned to the set of species common to all three data types per order. Distances were computed with the *phangorn* (Schleip et al., 2019) and *Quartet* (McGowan et al., 2025) packages in R.

Normalized RF and quartet distances were categorized as indicating high similarity (RF ≤ 0.25; quartet ≤ 0.10), moderate similarity (RF 0.25–0.50; quartet 0.10–0.30), substantial disagreement (RF 0.50–0.75; quartet 0.30–0.50), or very strong disagreement (RF > 0.75; quartet > 0.50). For Mycobacteriales, the high species richness precluded the reconstruction of 16S rRNA and WGS trees; phylogenetic signal and model inference for this order therefore rely solely on the timetree, and this limitation should be considered when interpreting its results.

For all downstream phylogenetic comparative modelling and statistical analysis, we adhered to the analytical framework described by Ajay et al. (2025). The backbone tree showing the phylogenetic relationships of the orders studied is given in Figure 1.

**Figure 1.**
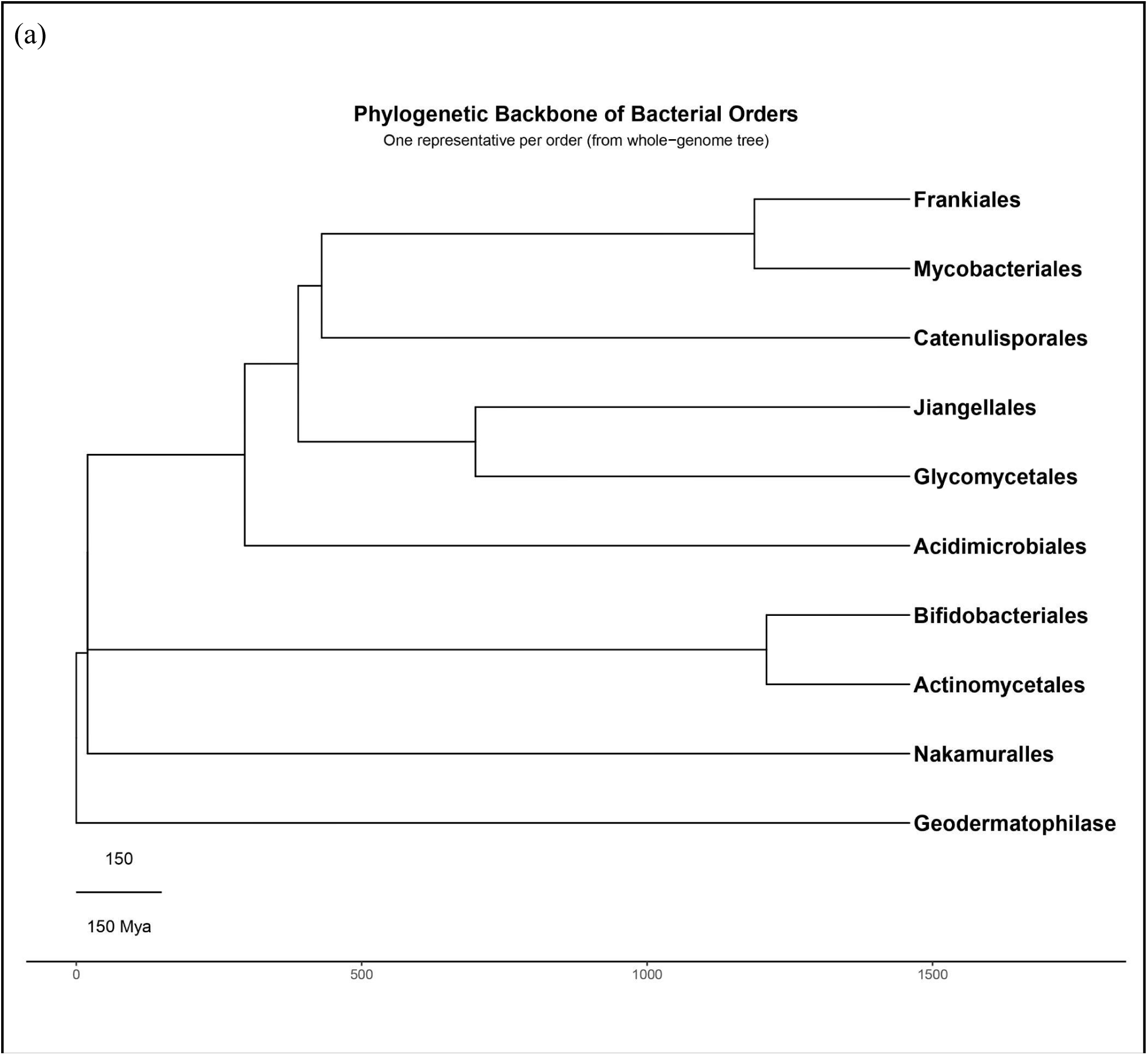
Phylogenetic framework for comparative analyses. Time–calibrated backbone tree of Actinomycetota used in this study. Tip labels indicate the actinobacterial orders included in the comparative analyses.

## 3. Results

Supplementary Table 2 (ST2) shows one-way ANOVA results and Fisher’s test for the distribution of genomic traits between the orders of the Actinomycetota. Actinomycetota exhibits inter-order heterogeneity in its genomic architecture, with all examined genomic traits showing significant differences in their distributions (P < 0.05). Mycobacteriales, Kineosporales and Glycomycetales showed a higher number of repeat regions and a greater repeat fraction in the genome as compared to the rest of the orders.

### 3.1 Discordant topologies: order–specific patterns and their interpretive implications

Topological congruence between time–calibrated, 16S, and whole–genome trees was assessed using normalized Robinson–Foulds (RF) and quartet distances (Table 2). For illustrative purposes, the timetree, 16S, and WGS phylogenies for Kineosporiales are shown in Figure 2; all tree files analysed in this study are deposited in the Zenodo repository [10.5281/zenodo.20495272]. Although the metrics operate on different scales (quartet range: 0.05–0.49; RF range: 0.17–0.89), their relative patterns were consistent. In *Actinomycetes*, the time and 16S trees showed moderate RF disagreement (0.415) but very high quartet similarity (0.052), indicating deep but localized split differences. Time versus WGS in the same order exhibited very strong RF discordance (0.887) and substantial quartet disagreement (0.492). *Bifidobacteriales* displayed substantial–to–very–strong discordance across all comparisons (RF: 0.537–0.805; quartet: 0.369–0.450). *Geodermatophilase* showed substantial RF disagreement (0.5–0.556) yet only moderate quartet discordance (0.214–0.381), consistent with few deep rearrangements. *Kineosporiales* demonstrated high similarity between time and WGS trees (RF = 0.167, quartet = 0.040), with moderate correspondence involving the 16S tree (RF = 0.333, quartet = 0.123). *Jiangellales* (n=5) was excluded from interpretation due to insufficient sample size. Thus, topological agreement is order–specific, and the choice of phylogenetic hypothesis markedly influences comparative outcomes.

**Table 2.**
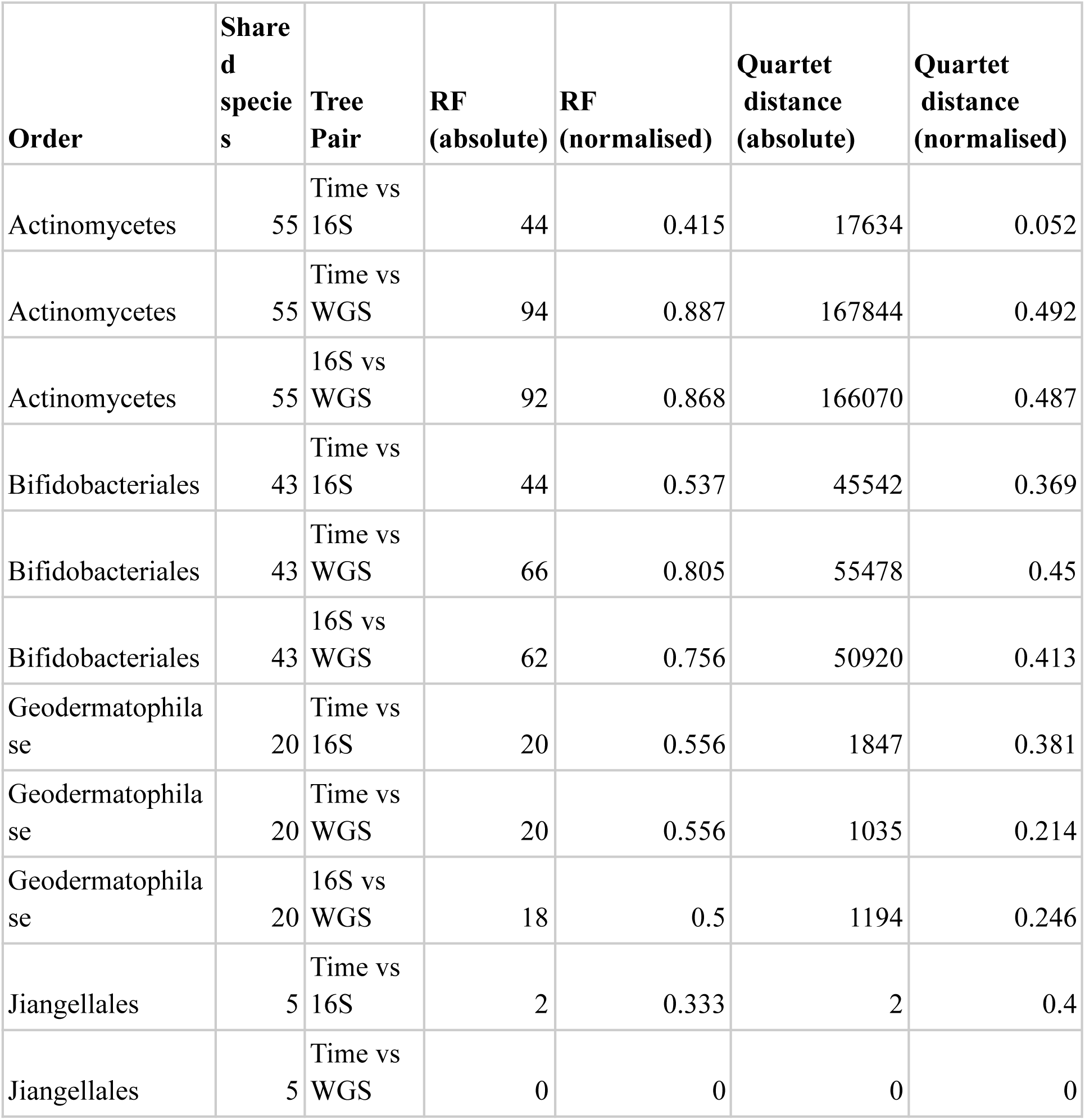
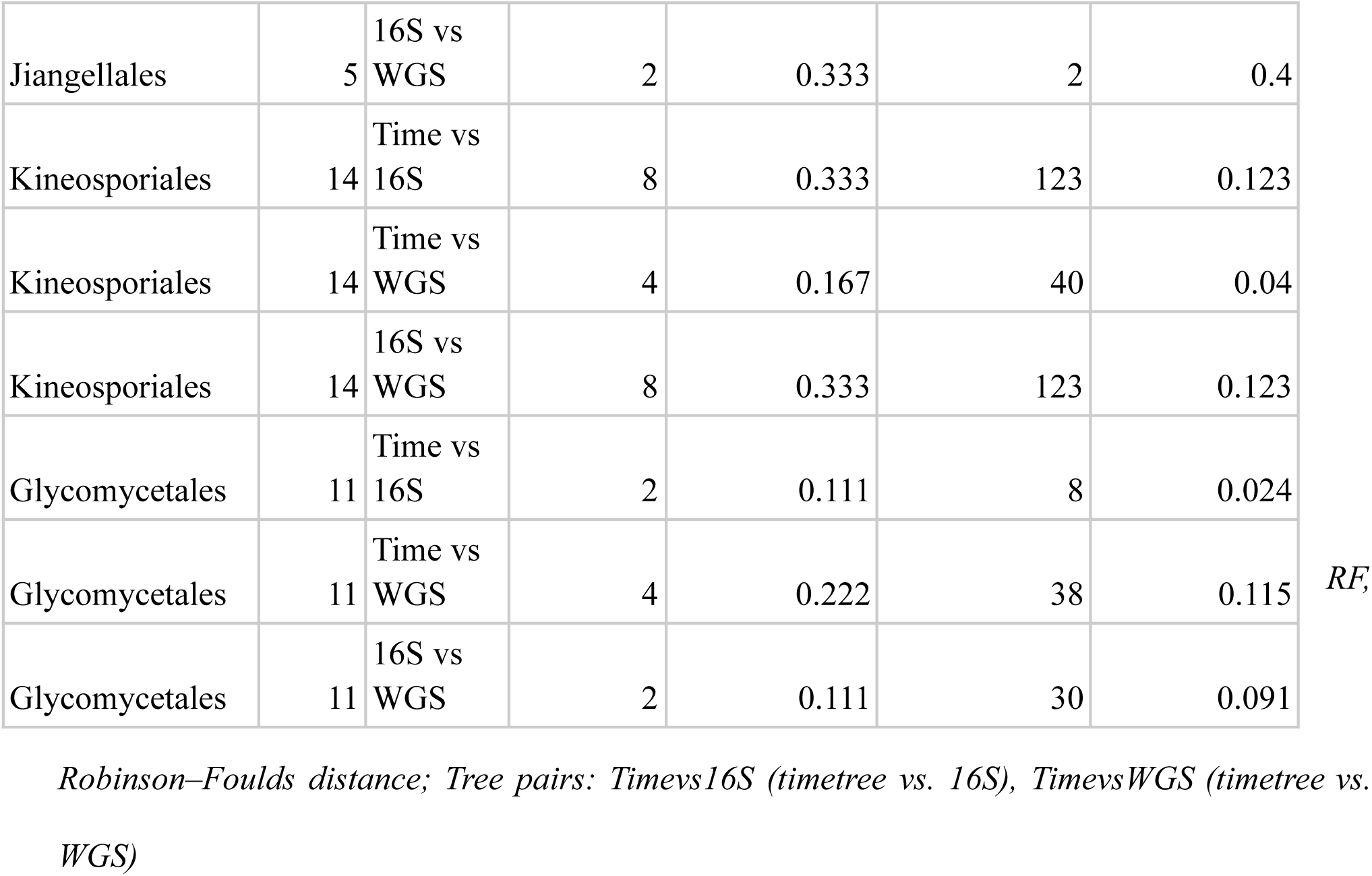
Topological congruence among the three phylogenies (timetree, 16S, and whole–genome). Normalised Robinson–Foulds (RF) distances and quartet distances are reported for each order and tree pair.

**Figure 2.**
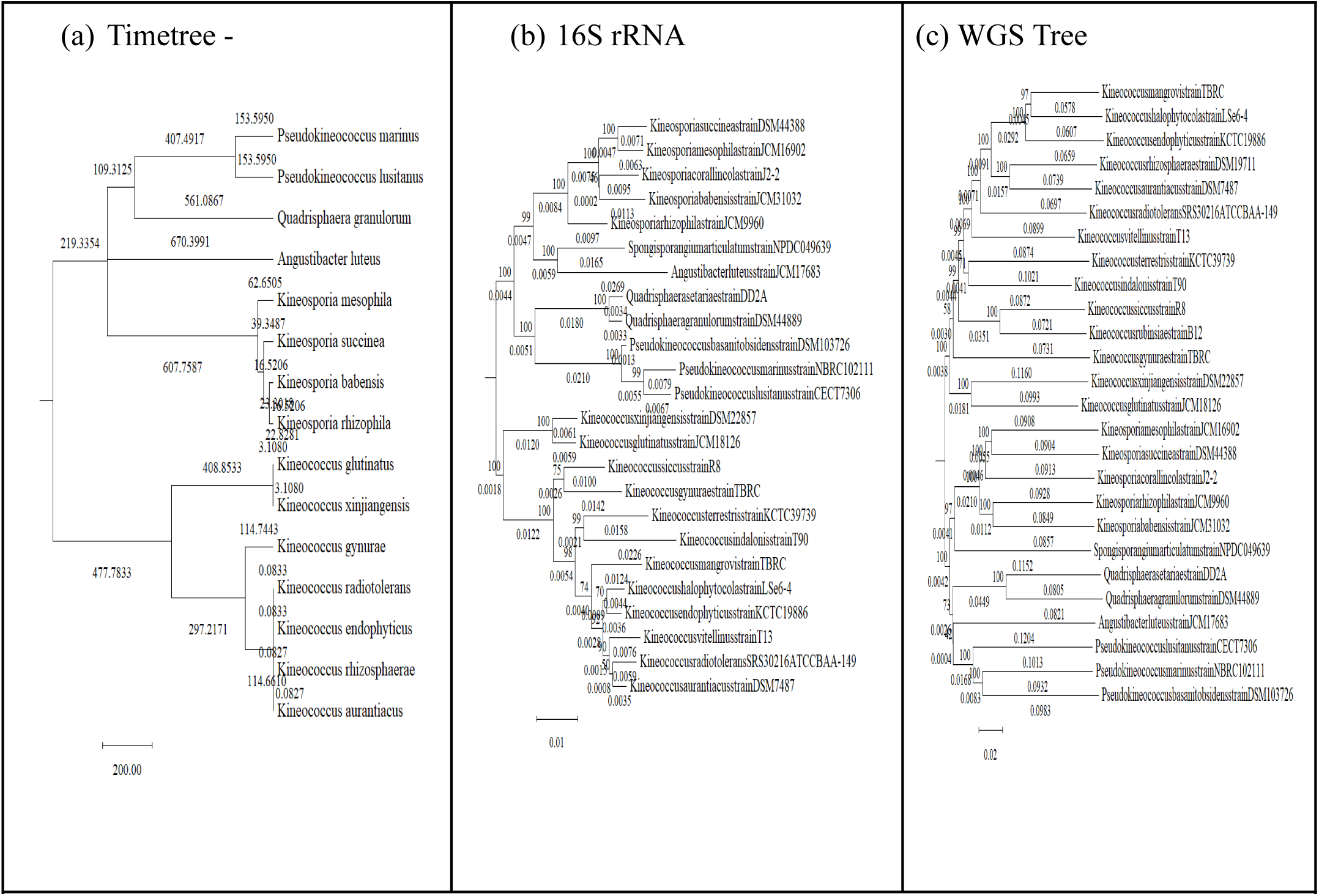
Representative phylogenies of Kineosporiales used in this study. (a) Timetree, (b) 16S rRNA gene tree, (c) WGS tree. All trees were pruned to match trait data availability. Full tree files are available on Zenodo [10.5281/zenodo.20495272]

### 3.2 Phylogenetic signal: conserved in core traits, plastic in repeats

Across all non–repeat genomic traits examined, 28 out of 50 trait–order comparisons showed complete agreement among the three phylogenies (16S, WGS, and timetree) regarding the presence or absence of a significant phylogenetic signal (Pagel’s λ) (Supplementary Table 3 (ST3)). The remaining 22 comparisons showed disagreement, consistent with the known sensitivity of phylogenetic signal metrics to differences in branch-length estimation and tree-reconstruction methods. Among non-repeating element traits, Genomic GC content exhibited a significant phylogenetic signal (Pagel’s λ) across all three phylogenies – 16S, WGS, and timetree – in every order examined. Gene count also showed a significant signal across all orders except Geodermatophilase, where the signal was consistently absent on all three trees. Across all three phylogenies, phylogenetic signal was consistently absent for average gene length (Actinomycetales), genome size and pseudogenes (Bifidobacteriales), most core traits (Geodermatophilase), and orientation genes (Glycomycetales).

Phylogenetic signal was not consistently detected across all three trees. Two major patterns were noted. First, traits that are genome–wide properties (e.g., number of protein-coding genes, repeat–containing genes, orientation bias) often showed significant signals in 16S and WGS trees but not in timetrees. This likely reflects the larger sample sizes in 16S/WGS phylogenies (e.g., 85 vs 45 species in Bifidobacteriales) and the fact that both marker–based and whole–genome similarity trees capture recent divergence more accurately than time–calibrated trees, which may compress branch lengths and reduce power. Conversely, traits such as average protein length and pseudogene counts in Actinomycetales, stress genes in Glycomycetales, and aquaporin genes in Bifidobacteriales were significant only on the timetree, suggesting that phylogenetic signal concentrated at deeper nodes may be better captured by time–calibrated branch lengths than by topologies scaled to substitution patterns.

Across all three phylogenies, repeat–related traits consistently lacked significant phylogenetic signal in most orders, with only three lineage– and topology–specific exceptions: a weak but significant signal in *Mycobacteriales* under the timetree (Pagel’s λ = 0.313, P = 0.004), significant Blomberg’s K in Bifidobacteriales under the WGS tree, and significant Blomberg’s K in *Kineosporiales* under the 16S tree. These exceptions indicate that while repeat content is generally evolutionarily plastic, lineage–specific phylogenetic structuring is detectable in these clades.

### 3.2 White-noise models, except for Bifidobacteriales, dominate repeat evolution

After testing for phylogenetic signal, we fitted evolutionary models to each trait using the timetree, 16S rRNA, and WGS phylogenies (Supplementary Table 4 (ST4)). The 16S and timetree topologies frequently agreed, both supporting a speciational (kappa) mode for gene counts and orientation traits in Actinomycetales and white noise for nearly all genomic traits in Geodermatophilus. The WGS tree often diverged, substituting EB for kappa in several Actinomycetales traits—a shift that does not change the biological interpretation, as both models indicate burst–like trait divergence. More importantly, the WGS tree recovered phylogenetic signal (BM, EB, delta) for traits that appeared as white noise on the 16S tree, including average gene length and aquaporin count in Bifidobacteriales and stress genes in Geodermatophilase, suggesting that genome–wide alignments may capture signals that single–marker trees miss.

Across all three phylogenies, repeat–related traits were predominantly best described by white noise, consistent with their largely plastic evolution (ST4). Three lineage–topology combinations departed from this pattern. Still, the inferred models reinforced the marginal nature of the detected signal: Mycobacteriales under the timetree remained white noise despite a weak yet significant λ, indicating negligible phylogenetic structure; Bifidobacteriales under the WGS tree was the sole case where repeat traits consistently favoured non–white models (a mixture of Brownian motion, early–burst, and delta processes across different repeat elements and repeat fraction), demonstrating that the whole–genome tree captures a genuine phylogenetic signature missed by the other trees; and *Kineosporiales* under the 16S tree supported an early–burst process, though this result must be interpreted cautiously given the order’s small sample size and the potential of single–marker trees to recover spurious structure. Overall, the WGS tree consistently uncovered phylogenetic signal in repeat traits that was not detected by the 16S or timetree analyses, underscoring its greater sensitivity for detecting evolutionary structure in rapidly evolving genomic features.

### 3.3 Repeat–genome interplay: lineage–specific composition, overall decoupling

We fitted phylogenetic generalized least–squares (PGLS) regressions to the timetree, 16S rRNA, and whole–genome (WGS) phylogenies to identify statistically significant (P < 0.05) pairwise associations between genomic repeat traits (total repeat length, GC–poor, and GC–rich repeats) and other core genomic features across the studied actinobacterial orders. The complete set of results is provided in Supplementary Table 5 (ST5), and significant associations are compiled separately in Supplementary Table (ST6). Across all three phylogenies, GC–rich repeat counts were positively associated with genome size and gene number in Mycobacteriales, Bifidobacteriales, and Actinomycetales, whereas GC–poor repeat counts were negatively correlated with genomic GC% in Mycobacteriales and Bifidobacteriales. In Mycobacteriales, despite a negative genome size–GC% relationship (R = −0.32, *P* < 0.001), total repeat content was more strongly tied to GC–rich (R = 5418, *P* < 0.001) than to GC–poor repeats, indicating that repeat expansion is driven primarily by GC–rich elements even in AT–enriched genomes. In Bifidobacteriales, GC–rich repeats similarly dominated genome size and gene–content associations. These patterns were robust to tree topology, confirming that the coupling between repeat composition and genome architecture reflects genuine evolutionary signal. Across most orders, the repeat fraction showed no significant phylogenetic association with core genomic traits, indicating decoupling of repeat–genome architecture. Exceptions were Bifidobacteriales, where repeat fraction correlated negatively with genomic GC% (WGS: R = -0.291, *P* = 0.03327, N = 49; 16S: R = -0.286, P = 0.04, N = 47), and Mycobacteriales (timetree), where repeat fraction was negatively associated with genome size, GC%, and protein–coding gene count, and positively with average protein and gene length (R values ranged from −0.498 to 0.440, all *P* < 0.05, N = 75). While genomic architecture influences the composition of repeat classes (GC–rich vs. GC–poor), the total quantity of repeats—as captured by repeat fraction—remains largely decoupled from core genomic features in most lineages.

Across all three phylogenies (timetree, 16S rRNA, and WGS), genome size scaled positively with both total gene count and protein–coding gene number in every order examined (e.g., R in Mycobacteriales: 0.94–0.97; Bifidobacteriales: 0.51–0.53; Geodermatophilase: 0.94–0.96; Glycomycetales: 0.98–1.00; all *P*<0.001), demonstrating that this fundamental genomic relationship is recovered consistently irrespective of the underlying tree topology. Across all three phylogenies, genome size was negatively correlated with genomic GC% in most orders—indicating that genome expansion is predominantly accompanied by the proliferation of AT–rich sequences—with positive associations observed only in Bifidobacteriales (timetree) and Glycomycetales (16S and WGS). In Bifidobacteriales, genomic GC% was positively associated with GC–rich repeat counts and negatively with GC–poor repeats (timetree: slopes −0.97, 1.15; both p<0.001), whereas in all other orders examined, genomic GC% showed no significant relationship with repeat content, indicating that base composition influences repeat–class proportions only in a lineage–specific manner.

### 3.4 Pulsed evolution in phylogenetically structured genomic traits

Across the timetree, 16S rRNA, and WGS phylogenies, the Anderson–Darling and Kolmogorov–Smirnov tests mostly rejected Brownian motion and Ornstein–Uhlenbeck expectations for traits that carried significant phylogenetic signal (excluding those best fitted by white noise) —a pattern indicative of pulsed evolution (Supplementary Tables 7, ST7). While genome size and GC% have previously been noted to evolve in a punctuated manner, our study recovers this signature across three independent topologies. It extends it to a wider suite of genomic features. In Mycobacteriales, Actinomycetales, and Bifidobacteriales, gene count, protein–coding gene count, average protein and gene length, stress genes, DNA–binding domains, pseudogenes, gene–orientation metrics, and aquaporin numbers all displayed significant λ and K and simultaneously rejected BM/OU (e.g., Mycobacteriales timetree: gene count AD = 15.05, *P* < 10⁻²⁴, KS = 0.138, *P* = 1.2 × 10⁻⁶; average gene length AD = 61.33, *P* < 10⁻²⁴, KS = 0.294, *P* = 1.3 × 10⁻²⁸). Geodermatophilase was a notable exception: several traits with robust phylogenetic signal (e.g., 16S Gc% λ = 0.99, *P* < 10⁻¹²; WGS genome size λ = 0.84, p = 0.019) did not reject BM/OU (AD-*P* = 0.057–0.14; KS *P* = 0.58–0.82), suggesting a lack of detectable punctuation in this order. In Bifidobacteriales, where the WGS tree uniquely captured significant phylogenetic signal in repeat traits (total length λ ≈ 1, *P* = 0.001; GC–poor repeats λ ≈ 1, *P* < 10⁻⁶), these repeats also rejected BM/OU (total number of repeats AD *P* < 10⁻²⁴, KS *P* < 10⁻¹⁹), indicating that pulsed repeat accumulation may have contributed to the punctuated pattern of genome–size evolution in this lineage. For Kineosporiales (n = 15) and Geodermatophilase (n = 20), the more parameter–rich PE2 and PE3 pulsed evolution models (and their BM variants) could not be reliably fitted due to limited sample size in timetrees and were therefore excluded from model comparisons. Across the three topologies, non–repeat genomic traits consistently reject gradual evolutionary models, revealing a pervasive signature of pulsed changes; repeat elements, by contrast, exhibit such departures only within the WGS–resolved Bifidobacteriales—a clade–specific exception not observed in any other orders. The full set of trait–distribution plots against the BM and OU expectations can be found in the accompanying Zenodo repository [10.5281/zenodo.20496329].

Re–evaluation with explicit pulsed models under a Burnham & Anderson ΔAIC framework confirmed that the distribution–based signals were not false positives for non–repeat traits (Supplementary Table 8 (ST8)). Mycobacteriales (timetree), Bifidobacteriales, Actinomycetales, and Geodermatophilase all showed strong support for pulsed evolution across the available topologies, with the timetree, 16S, and WGS topologies converging for the latter three orders. In all cases where pulsed evolution was favoured, the best–supported model was PE1_BM—a single pulse–size class operating against a continuous Brownian background (ΔAIC > 10). Kineosporiales, by contrast, showed no AIC support for evolutionary pulses in its non–repeat traits, although its small sample size warrants caution. *Bifidobacteriales* was the sole lineage in which repeat–related traits also achieved robust pulsed support: under the WGS tree, both GC–rich and GC–poor repeat counts strongly favoured pulsed evolution (PE1_BM; ΔAIC > 56 and > 192, respectively). Across the three phylogenies, support for pulsed evolution was qualitatively concordant, though the whole–genome tree consistently yielded the largest estimated jump magnitudes. Thus, while non–repeat genomic traits consistently exhibit phylogenetically structured, pulsed dynamics across multiple actinobacterial clades, in Bifidobacteriales, repeat elements uniquely mirrored the pulsed dynamics observed in non–repeat traits, a pattern not seen in any other order.

### 3.5 Genome size expansion is driven by gene content, not by repeat accumulation

Across the timetree, 16S rRNA, and whole–genome (WGS) phylogenies, gene content—quantified as total gene count and protein–coding gene count—was consistently the dominant predictor of genome size in every order examined (e.g., Mycobacteriales, Bifidobacteriales, Actinomycetales, Kineosporiales, Frankiales, Glycomycetales), with absolute coefficients an order of magnitude greater than those of any other trait (Supplementary Table 9 (ST9), Figure 3). This robust, topology–invariant relationship reinforces the long–standing view that genome expansion in bacteria is driven primarily by the accumulation of coding sequences. Genomic GC% emerged as a significant but secondary factor in several orders, including Actinomycetales (all three trees), Bifidobacteriales (timetree), Kineosporiales (all trees), and Frankiales (WGS); however, its effect size was invariably smaller than that of gene content, and its direction varied among lineages—positive in some, negative in others—indicating that base composition exerts a lineage–specific modulatory influence rather than a universal constraint on genome size. In sharp contrast, repeat–related traits (total repeat length, GC–poor and GC–rich repeat counts) were largely inconsequential for genome size evolution across the studied orders. The only repeat–trait significantly associated with genome size beyond Actinomycetales—GC–poor repeats in Bifidobacteriales under the 16S tree (*P* = 0.003)—was unsupported by the timetree and WGS trees. Given the known topology–sensitivity of repeat signals in this order, this isolated result likely reflects tree–specific covariance rather than a robust biological effect. For illustrative purposes, the phylogenetic regression coefficients for genome size as evaluated by timetree, 16S, and WGS phylogenies for Kineosporiales are shown in Figure 3; all tree files analysed in this study are deposited in the Zenodo repository [10.5281/zenodo.20495272].

**Figure 3.**
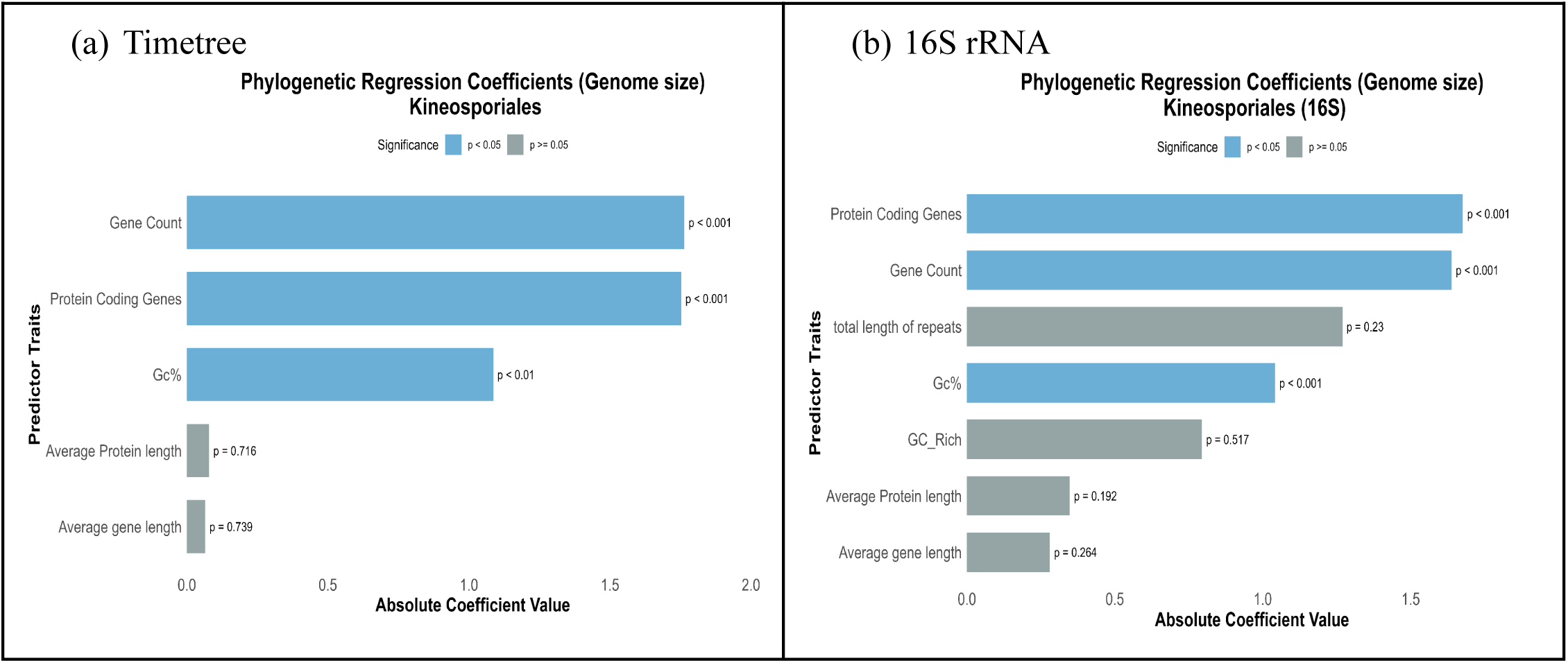
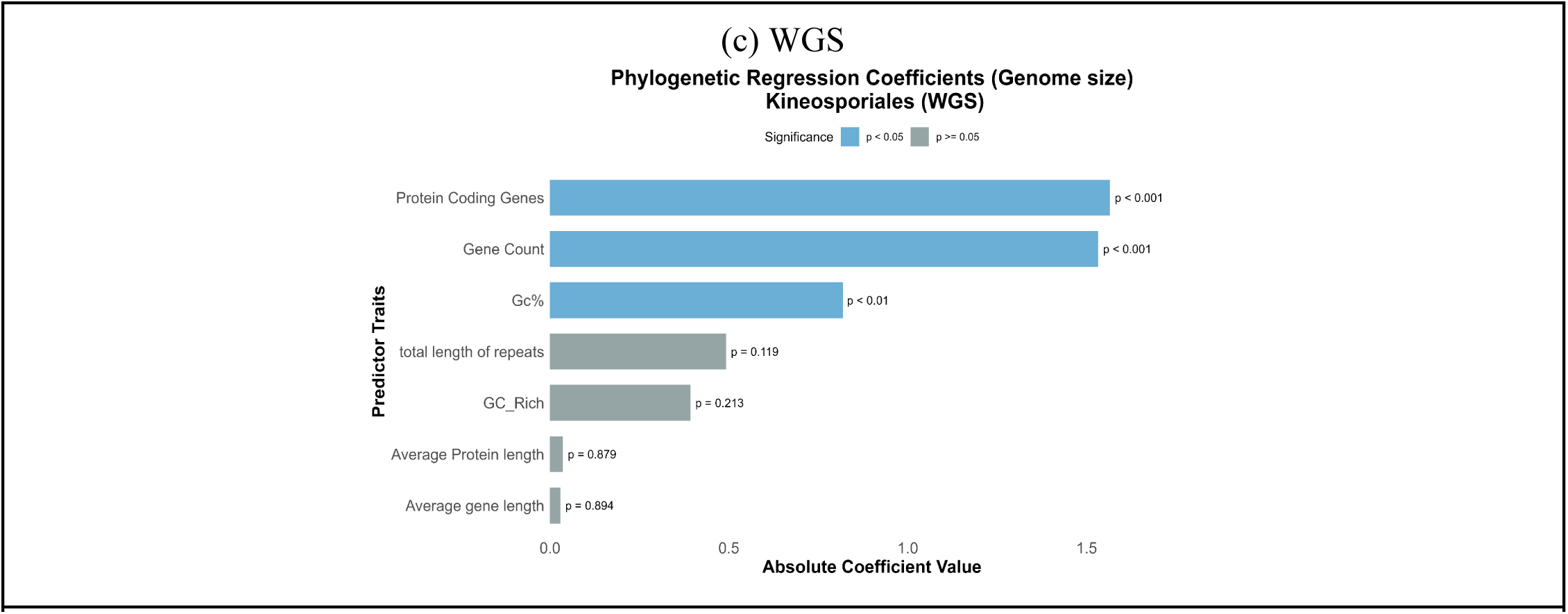
Phylogenetic regression coefficients for genome–size predictors in Kineosporiales, estimated via phylogenetic mixed models (pglmm) under the timetree, 16S rRNA, and whole–genome (WGS) topologies. Blue bars indicate significant predictors (P < 0.05); gray bars indicate non–significant predictors. Kineosporiales is shown as a representative example; full results for all orders are available in Supplementary Table 9 (ST 9)

## 4. Discussion

Unlike the 16S and WGS trees, the time–calibrated tree lacks strain–level resolution and does not include all sampled species, potentially missing variation captured by the other two topologies. While a growing body of work demonstrates that many organismal and genomic traits bear a clear phylogenetic signature and evolve in a pulsed, non–gradual manner—across both microbes (Gao and Wu, 2022) and arthropods (Ajay et al., 2025)—the present study reveals a strikingly different picture for genomic repeats in Actinomycetota. Irrespective of how they are quantified—whether as raw counts, as the fraction of the genome occupied by repeats, or partitioned into GC–rich and GC–poor classes—repeat–related traits consistently lack significant phylogenetic signal and show little to no evolutionary association with core genomic features such as genome size, gene number, or base composition. This decoupling holds across all three phylogenetic hypotheses examined (timetree, 16S rRNA, and whole–genome trees), indicating that the pattern is not an artefact of tree topology. The absence of signal is particularly notable given that repetitive elements have been shown to carry phylogenetic information in some plant and insect taxa, where trees reconstructed from repeat–marker distances can mirror those built from conventional markers (Dodsworth et al., 2015). While the utility of repeats as phylogenetic markers falls outside our scope, our results demonstrate that, as evolving traits, repeats behave largely independently of the genome–wide evolutionary forces that shape other genomic characteristics. Across the actinobacterial orders examined, repeat traits lack the phylogenetic influence and evolutionary coupling with core genomic features that characterise other genomic traits. Bifidobacteriales is the sole exception, and its signal appears only under the WGS tree.

This aligns with earlier findings in *Staphylococci* (Ajay et al., 2023), where repeat fraction lacked phylogenetic signal in contrast to other genomic traits. The current study extends that pattern to Actinomycetota, demonstrating that even a more refined repeat classification—by GC content and length class—reveals no signal, suggesting that such decoupling may be a general feature of bacterial genomes, pending broader taxonomic verification. Phylogenetic comparative analyses are sensitive to tree topology, yet across the three phylogenies employed here—timetree, 16S rRNA, and WGS—the broad patterns were notably congruent. This congruence likely reflects that topological differences tend to be concentrated in deeper nodes, where they exert less influence on trait covariance than changes near the tips, and that topological error generally erodes genuine phylogenetic signal rather than generating spurious structure (Symonds, 2002). Where the trees diverged, these discrepancies proved informative rather than contradictory—revealing, for example, the WGS–specific repeat signal in Bifidobacteriales—and highlighted the value of a multi–topology framework in capturing nuances that a single tree would miss. The detection of phylogenetic signal is also contingent on the taxonomic breadth under study (Martiny et al., 2013; Goberna & Verdú, 2016): at broad scales such as the order–level comparisons employed here, only traits with strong, deeply conserved phylogenetic structure are likely to emerge as significantly structured. It is well established that traits with developmental underpinnings tend to retain such signal, whereas those tied to ecological specialisation or niche colonisation are more evolutionarily labile (Goberna & Verdú, 2016; Ajay et al., 2025). Genomic repeats can fall into the latter category: they are known to contribute to adaptive responses to environmental stress in both plants (Sureshkumar et al., 2025) and bacteria (Zhou et al., 2014), and to generate variation within populations. Their generation via intrinsically stochastic slippage events (Tachida & Iizuka, 1992; Hancock, 1996) further supports the view that repeat–based traits are prone to evolutionary lability, lacking the phylogenetic covariance as seen in other genomic features. This lability may be reinforced by selection for compact genomes (Sela et al., 2016; Bobay & Ochman, 2017), amplified by large effective population sizes (Jordan et al., 2002), which likely promotes continuous purging of non–functional repeats while functional elements are dynamically generated (Delihas, 2011); with repeat landscapes ultimately shaped by ecological speciation processes in bacteria (Lassalle et al., 2015) . Whether the long–term temporal stability of repeat length distributions observed within a single species (McGinty et al., 2025) exists in bacteria remains an open question, but our cross–species analysis reveals no detectable phylogenetic imprint, underscoring the evolutionary lability of repeat–based traits.

Bifidobacteriales stands apart in its repeat landscape—showing a lower ratio of GC–rich to GC–poor repeats and emerging as an outlier in post–hoc comparisons. Its unique niche as a gut commensal or pathogen (Zhang et al., 2016), coupled with a specialised metabolism (Gupta et al., 2017) and high rates of horizontal gene transfer (Ventura et al., 2007), likely subjects it to coevolutionary pressures and repeat turnover not experienced by free–living actinobacteria. These factors may explain why repeat traits exhibit phylogenetic structure in this order alone. In *Bifidobacteriales*, WGS–based repeat traits share the early–burst and Brownian motion modes observed for GC%, gene count, and protein–coding gene count, and themselves exhibit pulsed evolution—PE1_BM being the favoured model—unlike repeat traits in any other order. These repeats are significantly associated with core genomic traits in PGLS regressions. Yet the WGS tree does not recover phylogenetic signal for genome size, a departure not only from the other orders and topologies examined here but also from the broadly conserved signal for genome size reported in other microbial and multicellular systems (Martiny et al., 2013; Gao and Wu, 2022). Gene count remains the primary driver of genome size in *Bifidobacteriales*, with repeats contributing a negligible role; a full account of trait interplay in this order may require a co–evolutionary perspective beyond the present scope.

Although genomic repeats were the primary focus of this study, our analyses also captured the broader evolutionary patterns of non–repeat genomic traits. Consistent with Martiny et al. (2013), who reported widespread phylogenetic signal in functional traits using 16S rRNA trees, we recovered similar signal across our timetree and WGS phylogenies, with early–burst, kappa, and delta models predominating. Extending the work of Gao et al. (2022), who demonstrated pulsed evolution for six genomic traits on a multi–locus timetree, we show that this pulsed signature is robust across alternative topologies—16S, WGS, and timetree—and applies to an expanded repertoire of genomic features, including gene length, protein length, stress gene number, coding gene count, gene orientation, and aquaporin count. For these traits, all of which carry significant phylogenetic signal (Martiny et al., 2013; Goberna & Verdú, 2016), the prevailing selection of kappa and early–burst models further indicates that much of their evolutionary change is concentrated around speciation events. Consistent with the established phylogenetic signal in most microbial genes (Comas et al., 2007) and gene–gene interactions (Balogun et al., 2025), gene count and protein–coding gene count emerge as the dominant drivers of genome size evolution, indicating that the pulsed dynamics of genome size primarily reflect concomitant pulses in gene content. GC composition exerts an additional, likely indirect, influence, mediated through the expansion of GC-rich protein–coding regions (Bohlin et al., 2017; Kyracaou et al., 2024). These patterns point to an interplay between selection and speciation in shaping genome architecture: with GC content elevated in core coding regions (Bohlin et al., 2017), pulses in the core genome may drive fundamental genomic rewiring, thereby facilitating actinobacterial diversification and niche adaptation. The consistent selection of PE1_BM as the best–supported pulsed model for non-repeat genomic traits across orders and all three phylogenies aligns with the pulsed–evolution patterns reported by Gao et al. (2022), and extends their finding from a single multi–locus tree to three independent topologies—timetree, 16S, and WGS—with genome size, protein–coding gene count, and total gene count exhibiting the highest jump rates (λ) across all clades.

The larger pulse magnitudes consistently recovered under the WGS tree likely reflect the strengths of the GBDP approach (Meier-Kolthoff et al., 2013; Meier-Kolthoff and Goker, 2019), which derives intergenomic distances from whole–genome features—including gene content, genome size, and base composition—rather than from a single marker (16S tree) or consensus calibration(timetree)(Kumar et al., 2017). GBDP captures both deep and recent divergence, making it particularly suited to resolving the phylogenetic structure of genome–level traits. In contrast, 16S rRNA trees, though widely employed as a phylogenetic standard, are constrained not only by substitution saturation at a single locus but also by intragenomic heterogeneity, poor concordance of hypervariable regions with core genome phylogenies (Hassler et al., 2022), and occasional horizontal gene transfer (Kitahara & Miyazaki, 2013). TimeTree timetrees average heterogeneous published divergence estimates, yet their resolution at shallow nodes is limited by sparse microbial fossil calibrations (Szöllősi et al., 2022), reliance on simple averaging of conflicting estimates (Kumar et al., 2022), and inflated relaxed–clock uncertainty at recent timescales (Battistuzzi et al., 2010). Greater phylogenetic resolution should, in principle, obscure pulsed signals, as the Brownian motion background more readily absorbs finer–scale variation. That pulse magnitudes are largest under the WGS tree, despite this, suggests that it captures genuine, speciation–associated genomic shifts rather than inflating jump estimates artefactually. The generally larger pulse magnitudes in species–rich orders such as *Mycobacteriales* (Herrera et al., 2025) and *Actinomycetales* (Barka et al., 2016) are consistent with earlier findings linking genomic pulse size to species richness (Ajay et al., 2025). *Geodermatophilase*, a smaller and sparsely sampled clade, deviates from this trend, and its unusually large pulse estimates recovered across three topologies must therefore be interpreted with caution. Timetrees do offer complementary insights to WGS-based ascomparative analyses assume that trait covariance scales with time, time–calibrated trees—in which branch lengths represent absolute divergence times—may be better suited for traits whose phylogenetic signal is concentrated at deeper nodes (Harvey & Pagel, 1991; Burbrink & Pyron, 2011).

A limitation of our study is that the time–calibrated tree lacks strain–level resolution and does not include all sampled species, potentially missing variation captured by the other two topologies and contributing to the observed differences in phylogenetic signal and evolutionary inferences across trees.

Across three independent phylogenetic hypotheses—timetree, 16S rRNA, and whole–genome trees—our study demonstrates that genomic repeats in Actinomycetota are evolutionarily plastic traits, largely decoupled from both phylogeny and the core genomic features that govern genome size. Genome expansion in these bacteria is driven primarily by the proliferation of protein–coding genes, a process likely tied to niche adaptation, while repeat accumulation follows an independent, lineage–specific trajectory. The present work focused on simple sequence repeats; extending this framework to additional repeat classes, such as transposable elements and CRISPR arrays, will be necessary to establish the generality of these findings. Whether repeats in other lineages—particularly the repeat–rich genomes of eukaryotes—exhibit phylogenetic signal remains an open question; resolving it will determine whether the decoupling observed here is a general feature of genome evolution or specific to prokaryotic systems.

## Acknowledgements

The research infrastructure used in this study was supported by a grant from the Department of Biotechnology (DBT), Government of India [Grant Number: BT/PR59599/BID/7/1087/2025].

## Declaration of AI-Assisted Writing Support

The authors acknowledge the use of generative AI for language polishing, grammatical correction, and sentence refinement. The authors solely conducted all scientific content, data analysis, interpretation, and conceptual development. The AI served only as a writing assistant and did not contribute to any scientific reasoning or insights. The final manuscript has been reviewed and approved by all authors, who take full responsibility for its scientific integrity.

## Author Contributions

Chandana Gobattini and Uday Kumar Ammisetty contributed equally to this work. Chandana Gobattini: Data Curation (genomic data collection), Investigation (tandem repeat data generation using Tandem Repeat Finder), Visualisation (ANOVA plots). Uday Kumar Ammisetty: Data Curation (genomic data collection), Investigation (tandem repeat data generation using Tandem Repeat Finder). Bhargav Reddy Konkala: Investigation (repeat data generation). Akash Ajay: Conceptualisation, Methodology, Formal Analysis, Writing – Original Draft. All authors reviewed and approved the final manuscript.

## Data Availability Statement

All data and R scripts used in this study, including morphological and genomic datasets and analysis code for phylogenetic and random forest models, will be made publicly available upon acceptance of the manuscript. The data and scripts will be deposited in a public GitHub repository to ensure accessibility for further research and replication.

